# iEDGE: integration of Epi-DNA and Gene Expression and applications to the discovery of somatic copy number-associated drivers in cancer

**DOI:** 10.1101/573824

**Authors:** Amy Li, Bjoern Chapuy, Xaralabos Varelas, Paola Sebastiani, Stefano Monti

**Affiliations:** Division of Computational Biomedicine, Boston University School of Medicine, Boston, MA 02118, USA; Bioinformatics Program, Boston University, Boston, MA 02215, USA; Dana-Farber Cancer Institute, Department of Medical Oncology, Brookline, MA 02215, USA; University Medical Center Göttingen, 37075 Göttingen, Germany; Department of Biochemistry, Boston University School of Medicine, Boston, MA 02118, USA; Department of Biostatistics, Boston University School of Public Health, Boston, MA 02118, USA

**Keywords:** Cancer Genomics, Integrative Analysis

## Abstract

The emergence of large-scale multi-omics data warrants method development for data integration. Genomic studies from cancer patients have identified epigenetic and genetic regulators – such as methylation marks, somatic mutations, and somatic copy number alterations (SCNAs), among others – as predictive features of cancer outcome. However, identification of “driver genes” associated with a given alteration remains a challenge. To this end, we developed a computational tool, iEDGE, to model cis and trans effects of (epi-)DNA alterations and identify potential cis driver genes, where cis and trans genes denote those genes falling within and outside the genomic boundaries of a given (epi-)genetic alteration, respectively.

First, iEDGE identifies the cis and trans genes associated with the presence/absence of a particular epi-DNA alteration across samples. Tests of statistical mediation are then performed to determine the cis genes predictive of the trans gene expression. Finally, cis and trans effects are annotated by pathway enrichment analysis to gain insights into the underlying regulatory networks.

We used iEDGE to perform integrative analysis of SCNAs and gene expression data from breast cancer and 18 additional cancer types included in The Cancer Genome Atlas (TCGA). Notably, cis gene drivers identified by iEDGE were found to be significantly enriched for known driver genes from multiple compendia of validated oncogenes and tumor suppressors, suggesting that the remainder are of equal importance. Furthermore, predicted drivers were enriched for functionally relevant cancer genes with amplification-driven dependencies, which are of potential prognostic and therapeutic value. All the analyses results are accessible at https://montilab.bu.edu/iEDGE.

## Introduction

A central goal of cancer genomics is to identify key genetic and epigenetic alterations that promote initiation and/or progression of cancer. These alterations can manifest themselves across multiple biological levels – DNA, RNA, protein – and can be correspondingly quantified by multiple high-throughput profiling technologies.

Large-scale cancer genomics data compendia such as The Cancer Genome Atlas (TCGA) have generated comprehensive multi-omics datasets for tens of thousands of patients across ∼30 types of cancer (Cancer Genome Research Network, 2013). The availability of such large-scale datasets provides an opportunity to develop methods that integrate data from multiple types of profiling platforms, supporting the discovery of novel candidate driver genes and eventually diagnostic and prognostic biomarkers and therapeutic targets.

Past research using integrative approaches have shown success in discovering novel cancer drivers (Akavia et al. 2010; Xie et al. 2012). However, many existing methods often rely on ad-hoc, albeit sophisticated, analysis methods and scripts not accessible to analysts other than those responsible for their development. Furthermore, the generated analysis results are often static, and not accessible in an interactive fashion. The approach presented here aims to address both of these shortcomings.

The central hypothesis behind integrative approaches is that the integration of multi-level genomics data allows for prioritization of putative cancer “drivers” that are potential biomarkers or therapeutic targets. An important genetic alteration type in cancer is somatic copy-number alterations (SCNAs). SCNAs harbor many known cancer drivers (oncogenes or tumor suppressors) and play an important role in cancer initiation and/or progression through activation of oncogenes and inactivation of tumor suppressors (Beroukhim et al. 2010; Zack et al. 2013). Identification of unknown SCNA-associated cancer drivers is complicated by the fact that each SCNA contains many genes, often even a complete chromosome arm, the majority of which is likely not to confer any selective advantage (i.e., passengers). One approach to address this problem is to prune the set of candidate drivers based on their association with paired omics data, such as gene expression profiles. For example, one can prioritize genes found in frequent SCNA peaks whose gene expression changes are associated with corresponding copy number changes (Monti et al. 2012, Lai et al. 2017). Even after this pruning step, the set of remaining candidate drivers might still yield too many testable hypotheses for use in functional validation studies. More importantly, association between SCNA and gene expression alone may not be the best metric for ranking potential cancer drivers.

To address this problem, we developed a methodology that identifies SCNA-(or other alteration-)associated genes and performs prediction of cis gene drivers prioritized by their capability of transactivating downstream gene expression (trans effects), by leveraging the trans-gene signature (gene sets outside the SCNA of interest) whose expression is also associated with a given SCNA event of interest. The approach is predicated on the hypothesis that SCNA-related drivers of tumorigenesis will mediate a larger proportion of the downstream effect observable by trans gene expression than non-drivers. Using this heuristic, and based on the Sobel test of mediation (Sobel 1982), we identify putative SCNA-associated cancer drivers as the cis gene mediating the most trans gene expression, although the method allows users to customize the set of trans genes considered for the ranking of putative cis gene drivers.

We developed a corresponding software tool, *integration of Epi-DNA and Gene Expression* (iEDGE), available at https://github.com/montilab/iEDGE, for the prediction of (epi-)DNA-associated cancer drivers. Some of iEDGE’s methodological components were previously applied to the cis/trans analysis of diffuse large B cell lymphoma multi-omics datasets, and have shown success in uncovering SCNA-associated cis driver genes (Chapuy et al. 2018, Monti et al. 2012). Here, we present an expanded and optimized method crucially incorporating the mediation step, and we present the results of its application to the prediction of SCNA-associated cancer drivers across 19 cancer types using data from TCGA, with a particular focus on analysis of TCGA breast cancer. Our list of candidate drivers is highly enriched for known oncogenes and tumor suppressors and additionally implicates many suspected drivers as well as novel candidate genes with potential prognostic or therapeutic importance in cancer.

## Results

### iEDGE identifies SCNA-associated cis and trans genes and pathway signatures in TCGA breast cancer

To identify cis and trans gene signatures of SCNAs in breast cancer, we performed integrative analysis on paired copy number and gene expression data from TCGA breast cancer primary tumors from four gene expression-based molecular subtypes (Luminal A, Luminal B, Her2 and Basal) using the workflow summarized in Figure 1.

**Figure 1.**
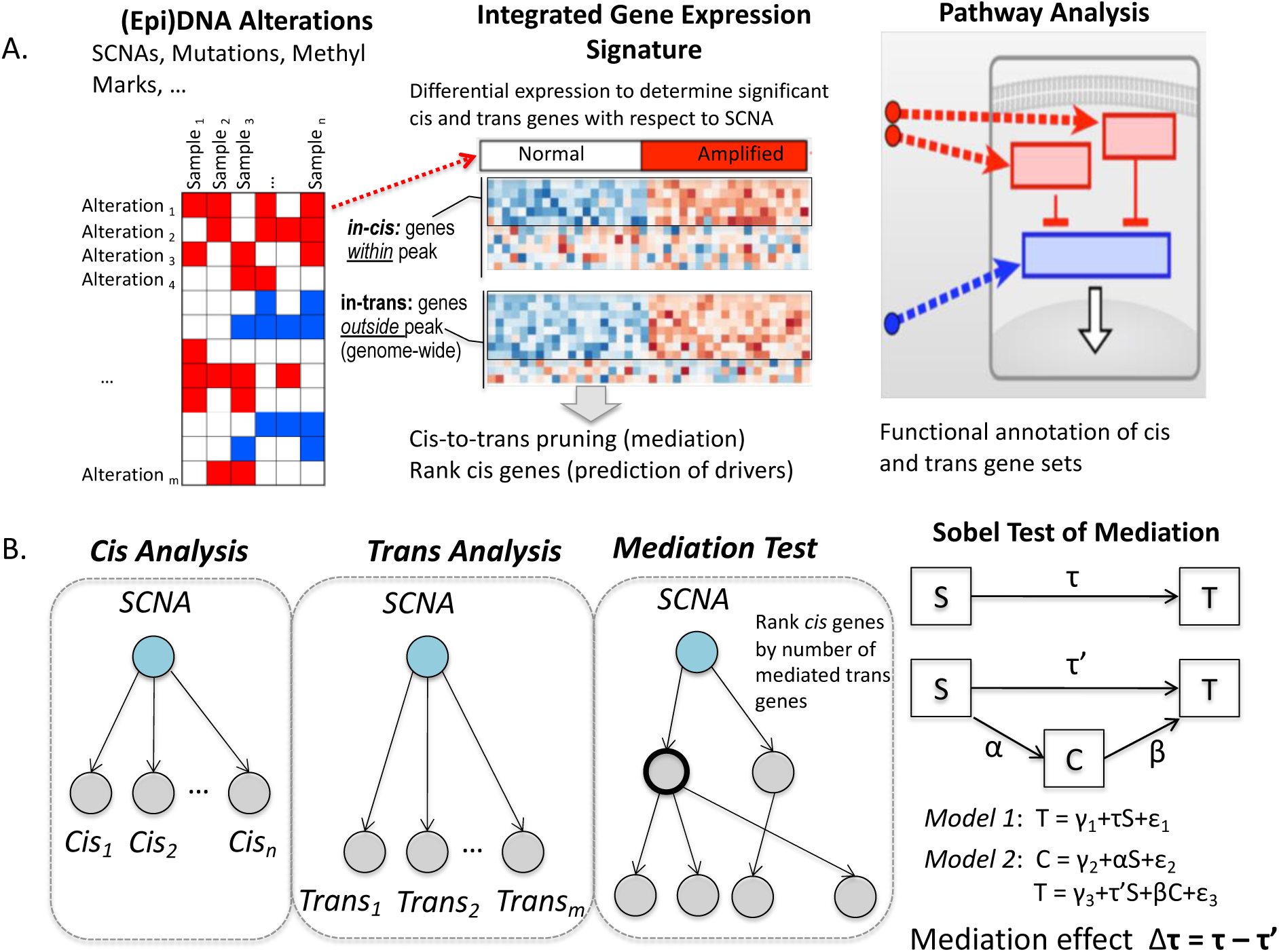
Overview of iEDGE workflow. (A) iEDGE identifies the cis and trans gene expression signatures of epi-DNA alterations, e.g., Somatic Copy Number Alterations (SCNAs), through differential expression analysis. It then performs pathway enrichment analysis to identify pathways or genesets associated with each SCNA. (B) Secondly, it performs cis-to-trans gene mediation analysis to identify putative driver cis genes of each epi-DNA alteration, defined as the cis genes that mediate the most trans gene expression.

Focal SCNAs were identified using GISTIC2.0 (Mermel et al., 2011), including 29 amplifications and 40 deletions. Next, we identified sets of cis and trans genes with significant differential expression with respect to each SCNA, and performed, pathway enrichments of cis and trans gene sets (Figure 1A).

A list of cis genes whose expression is significantly associated with each SCNA is summarized in Supplemental Table S1 (Fold change > 1.2, one-directional FDR <0.25). This list comprises an average of 20 genes (and a median of 8 genes) per SCNA, with 1330 genes in total (269 in amplifications, 1061 in deletions) out of the original 2003 genes across all SCNAs identified using GISTIC2.0, a number clearly still too large for meaningful consideration for functional validation.

Similarly, a list of significantly differentially expressed trans genes is summarized in Supplemental Table S2 (Fold change > 1.5, bi-directional FDR < 0.1). This list contains an average of 865 (a median of 598) significant trans genes per SCNA.

Pathway enrichment analyses of the union of cis and trans genesets yielded interesting and potentially biological meaningful patterns (Figure 2). In particular, several pathways were enriched across most SCNAs. For instance, gene sets HALLMARK_ESTROGEN_RESPONSE_LATE, HALLMARK_ESTROGEN_RESPONSE_EARLY, HALLMARK_G2M_CHECKPOINT were significant hits in more than 75% of SCNAs. These gene sets indicate global patterns of downstream effects related to genomic instability induced by co-occurring SCNAs across tumor samples (Figure S1), since co-occurring SCNAs will tend to have overlapping trans signatures, hence common pathway enrichments. To elucidate breast cancer subtype specific pathway enrichments, we categorized each SCNA by their enrichment in each of four major breast cancer types: Luminal A, Luminal B, Her2, Basal. In addition, for each pathway, we tested if the enrichment across SCNAs tended to occur in subtype-specific SCNAs compared to non-subtype-specific SCNAs (Figure 2). Several significant pathways were found to be occurring more frequently in Basal-specific SCNAs, including HALLMARK_SPERMATOGENESIS, HALLMARK_KRAS_SIGNALING_UP, HALLMARK_E2F_TARGETS, HALLMARK_BILE_ACID_METABOLISM, HALLMARK_FATTY_ACID_METABOLISM, HALLMARK_UV_RESPONSE_UP, HALLMARK_MYOGENESIS (FDR < 0.05). The enrichment of KRAS signaling can be explained by published evidence supporting KRAS activation in basal-type breast cancer cells compared to luminal cells (Kim et al, 2015).

**Figure 2.**
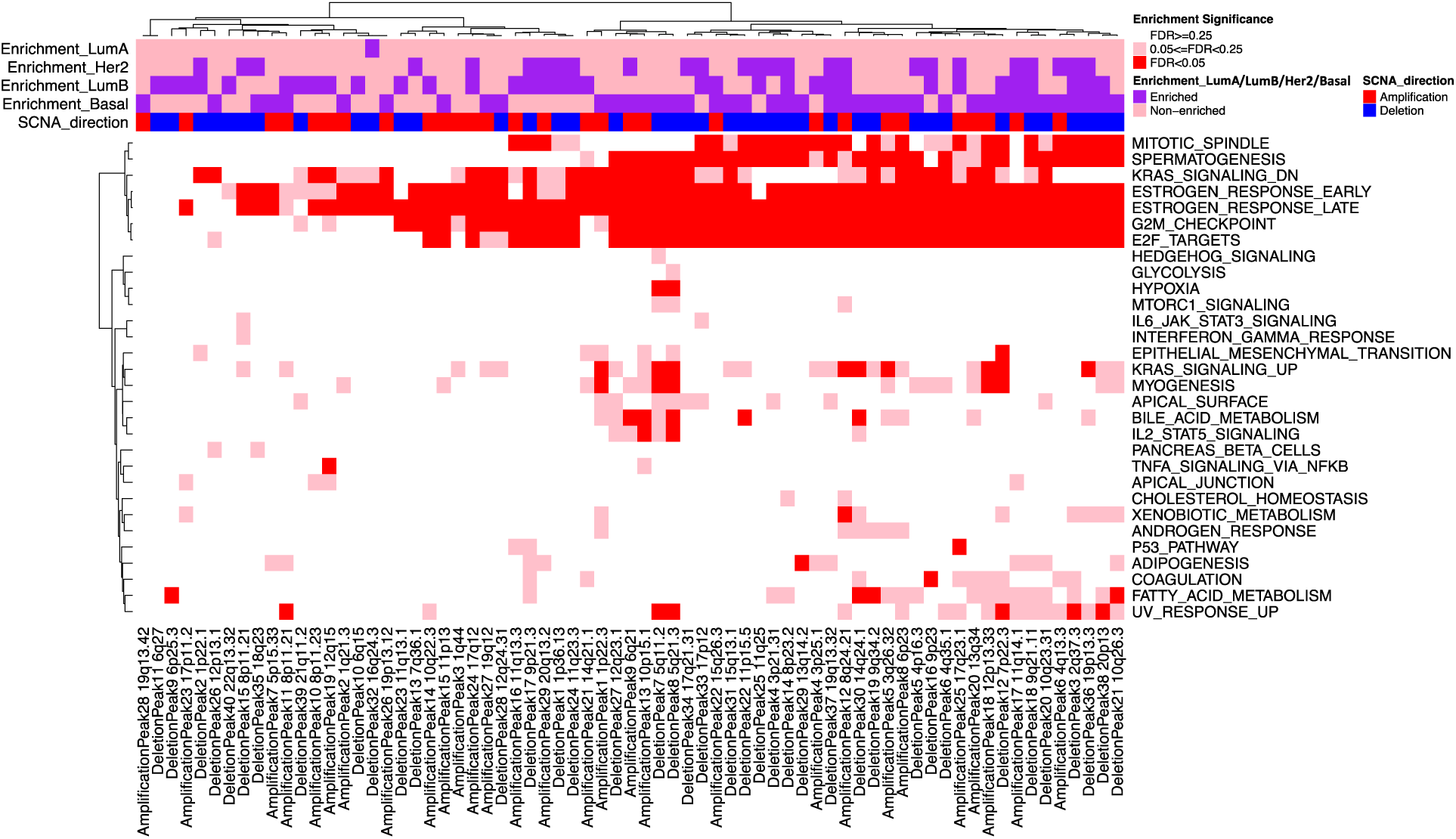
Pathway enrichment significance of cis and trans gene signatures of TCGA breast cancer somatic copy number alterations. Heatmap representing the enrichment of each SCNA-associated cis/trans union signatures (columns) with respect to the genesets (rows) in the MSigDB “Hallmark” compendium for the TCGA breast cancer dataset. Color-coded bars (pink/purple) on top of the heatmap indicate the enrichment of each SCNA in a particular breast cancer subtype as determined by Fisher exact test of the within-vs. outside-subtype counts of SCNA occurrences. The directionality of each SCNA is also represented as amplification (red) or deletion (blue).

### iEDGE identifies known cancer drivers in TCGA breast cancer

After the identification of the cis and trans gene signatures of SCNAs, iEDGE can be used to predict cis gene drivers of each SCNA (Figure 1B). For each SCNA in the TCGA breast cancer dataset, we ranked the significant cis genes using the Sobel test of mediation (Sobel 1982) to predict the driver gene of the alteration. Cis genes were ranked by the weighted fraction of significant trans genes they mediate (see Methods, and Supplemental Table S3). Rank-1 cis genes – the genes mediating the largest proportion of trans genes within each alteration, hence hypothesized to be driver genes for each SCNA – are summarized in Supplemental Table S4.

To verify the functional relevance of Rank-1 driver gene predictions, we tested the list of Rank-1 cis genes for enrichment in cancer driver databases compared to non-Rank-1 cis genes (Supplemental Table S5). Significance of enrichment was determined using a one-sided Fisher exact test on the contingency table of counts of Rank-1 *vs.* non-Rank-1 cis genes against the list of known cancer drivers *vs.* unknown genes. Predicted Rank-1 driver genes in amplifications were significantly enriched for known oncogenes in UNIPROT (P-value = 0.0016), COSMIC (P-value = 0.0012) and the combined test (union of driver databases) (P-value = 0.0041), and predicted Rank-1 driver genes in deletions were significantly enriched for tumor suppressors in TUSON (P-value = 0.0045), UNIPROT (P-value = 0.016), COSMIC (P-value = 0.002), and the combined test (P-value = 0.013). Among the 65 Rank-1 cis genes (Supplemental Table S4), known cancer drivers included *MCL1, ACTL6A, MYC, CCND1, FOXA1, ERBB2, CCNE1* (oncogenes in amplifications) and *RPL5, ZMYND10, KMT2C, CSMD1, CDKN2B, PTEN, CREBL2, FANCA* and *ARHGAP35* (tumor suppressors in deletions) (Supplemental Table S5).

### iEDGE identifies amplification-driven gene dependencies in TCGA breast cancer

In addition to the prediction of known cancer drivers, we assessed whether iEDGE was capable of identifying genes with copy-number driven cancer dependencies, specifically, amplification-driven gene dependencies (see Methods). To this end, we identified genes with increased essentiality in an amplified state using DepMap data of genetic screens (RNAi screens) paired with copy number data (CCLE) (Supplemental Table S6), and tested for their enrichment among iEDGE Rank-1 cis genes. In the TCGA breast cancer dataset, we found a highly significant enrichment of Rank-1 cis genes among genes with amplification-driven gene dependencies (Fisher test one-sided P-value = 3.53e-5). These were *ERBB2, CCNE1, CCND1, FOXA1, ANKRD17, MCL1*. All of these genes are linked to breast cancer development in the overexpressed state. *ERBB2, CCND1, CCNE1* are well-characterized oncogenes present among our curated set of cancer driver databases (see Methods). *FOXA1* has been shown to play an important role in promoting ER+ breast cancer (Meyer and Carroll 2012). *ANKRD17* is a cyclin E/Cdk2 substrate which positively regulates cell cycle progression by promoting G_1_/S transition (Deng et al. 2009). *MCL1* high expression is linked to poor prognosis in triple-negative breast cancer and targeting of *MCL1* restricts the growth of triple negative breast cancer xenografts, suggesting its potential therapeutic value (Campbell et al. 2018).

In summary, using the cis gene mediation step, we identified known SCNA-associated breast cancer gene drivers and novel genes with amplification-driven gene dependency that are of potential prognostic or therapeutic value.

### TCGA pan-cancer analysis

Next, the driver prediction procedure was carried out across 19 cancer types from TCGA (Supplemental Table S7). We tested for the enrichment of Rank-1 cis genes with known cancer drivers across the 19 cancer types and found significant enrichment (FDR < 0.05) in 15 out of 19 cancer types (Kolmogorov-Smirnov test of p-values against uniform [0,1]: p-value = 2.5e-13) (Supplemental Table S8).

Additionally, we tested for the enrichment of Rank-1 cis genes with genes manifesting amplification-driven dependencies and found significant enrichment (FDR < 0.05) in 8 out of the 19 cancer types analyzed, including BLCA, BRCA, CESC, ESCA, HNSC, LUAD, OV, UCEC (Kolmogorov-Smirnov test of p-values against uniform [0,1]: p-value = 0.00287) (Supplemental Table S9).

To assess the importance of the cis mediating step for the identification of known cancer drivers, we ordered Rank-1 cis genes (Supplemental Table S10) by the number of cancer types they occur in, and tracked which of these genes were validated cancer drivers. The derived ordered Rank-1 gene list was then compared with the ordered list of recurrent top differentially expressed (D.E.) cis genes in each SCNA, irrespective of their cis-mediating rank (Supplemental Table S11), as well as with the ordered list of *all* recurrent differentially expressed cis genes in each SCNA (Supplemental Table S12). These comparisons confirmed that the recurrent Rank-1 cis genes were more likely to capture known drivers than both the recurrent top D.E. cis genes and all cis genes (Figure 3A). Remarkably, among the top 15 Rank-1 cis genes, 14 genes were known cancer drivers (*PTEN, WWOX, CCNE1, CDKN2A, MAP2K4, EGFR, KAT6A, MYC, KRAS, ERBB2, WHSC1L1, PARK2, CREBBP, RB1*) and only 1 was unverified (*ATP9B*) (Figure 3B, left) (Fisher’s exact test P-value = 7.76e-08). In contrast, in the absence of the mediation step, among the top 15 top D.E. genes in SCNAs, 9 were confirmed drivers (Figure 3B, middle) (Fisher’s exact test P-value = 0.0047), and among the top 15 cis genes in SCNAs, only 4 were confirmed drivers (Figure 3B, right) (Fisher’s exact test P-value = 0.21). These results demonstrate the usefulness of the mediation test in further restricting the list of SCNA-associated cis genes to those of functional relevance across multiple cancer types. The mediation test provides a substantial improvement over ranking solely based on the cis gene differential expression and demonstrates the added value of performing integrative analysis that models downstream biological effects as captured by trans gene expression.

**Figure 3.**
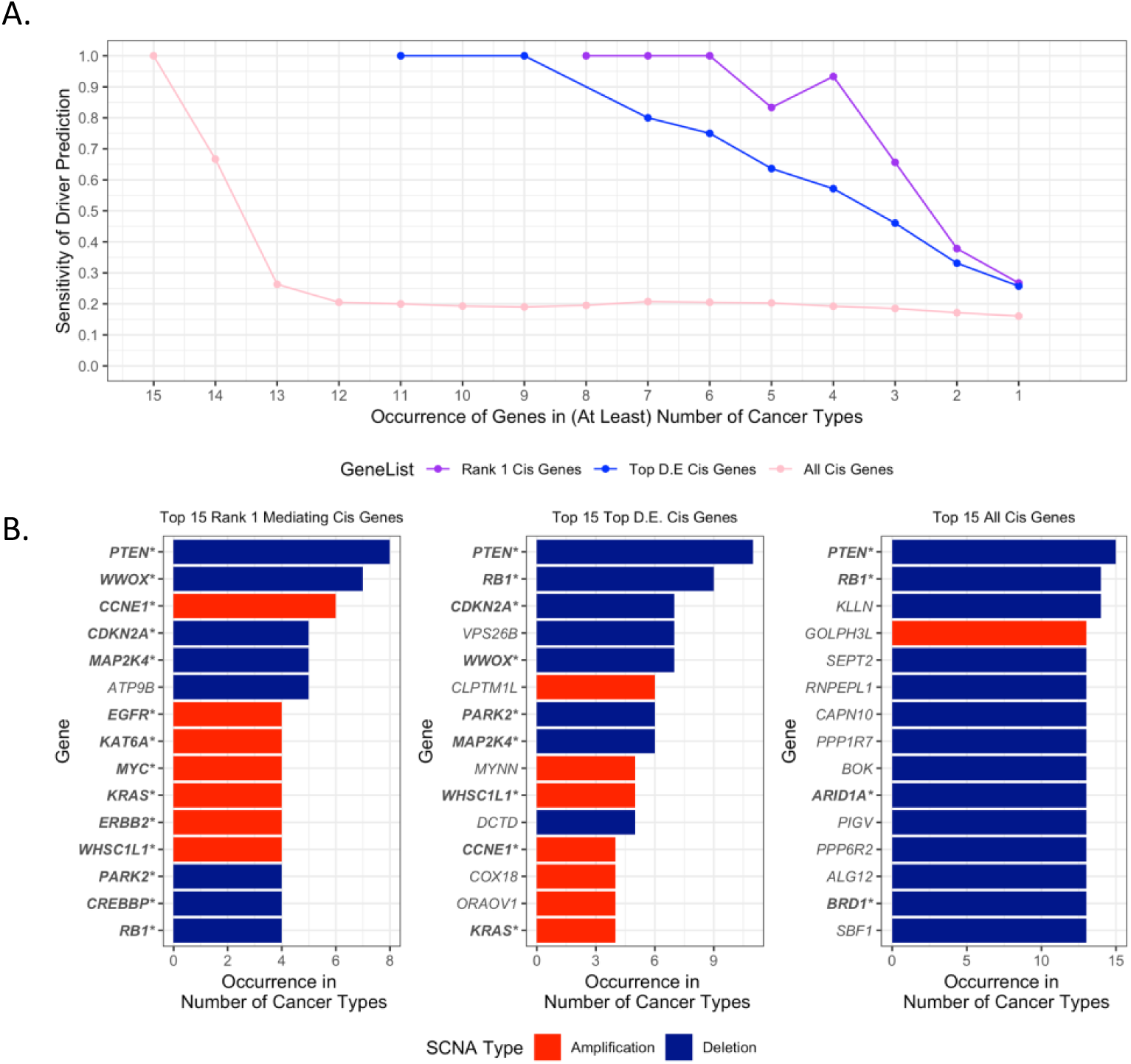
Sensitivity of cancer driver predictions across multiple cancer types. (A) Sensitivity of Driver Predictions (expressed as the fraction of known drivers in the ranked list of genes) vs. Occurrence of Genes in Number of Cancer Types (B) Barplots of top 15 genes ranked by occurrence in number of cancer types. Left: Rank-1 mediating cis genes; middle: top differentially expressed (D.E.) cis genes, right: all cis genes differentially expressed in alteration. Known cancer drivers are in bold and marked with (*).

In addition to enrichment for known cancer drivers, Rank-1 cis genes that frequently occur across cancer types (Supplemental Table S10) include putative or novel genes implicated in cancer initiation or progression. These include: *TRIP13*, a mitosis regulator that was shown to promote tumor growth in colorectal cancer (Sheng et al, 2018) and is a predictor of poor prognosis in prostate cancer (Dong et al. 2019); *ORAOV1*, a gene overexpressed in many solid tumors that is linked to generation of reactive oxygen species (Zhai et al. 2014); *TPX2*, an interactor and substrate of Aurora-A that is a potent oncogene amplified in many cancers and a promising therapeutic target (Kufer et al. 2002; Yan et al, 2016); and *DUSP22*, which has been shown to behave as a tumor suppressor gene in peripheral T-cell lymphomas (Mélard et al, 2016) and regulates ERα dependent transcription in breast cancer cells (Sekine et al, 2007).

### Evaluation of reproducibility of Rank-1 cis genes

To evaluate the reproducibility of cis gene ranking based on mediation testing, we quantified the consistency of Rank-1 cis gene predictions across bootstrapped resamples of the original TCGA breast cancer dataset (Figure 4). The majority of Rank-1 cis genes (69.2%) was consistently found as Rank-1 cis genes in bootstrapped datasets with greater than 0.75 fraction of inclusion, i.e., found as Rank-1 cis genes in 75% of the resampled datasets, and the fraction of inclusion is not biased by SCNA type (amplification or deletion) (Figure 4A). We further observed that the fraction of inclusion is positively associated with the Weighted Fraction of Trans Mediation (WFTM), the score used to rank cis genes within each SCNA based on the extent of trans genes mediation (see Methods: Mediation testing and prediction of cis drivers for calculation of WFTM) (Figure 4B). In particular, while Rank-1 cis genes with lower fraction of inclusion across bootstrapped resamples tend to have lower WFTM, Rank-1 cis genes with higher fractions of inclusion show more variability in WFTM but generally tend to have higher WFTM. The fraction of inclusion is negatively associated with the entropy of WFTM of cis genes in SCNAs of interest (Figure 4C). This is an indication that SCNAs with a single dominant cis gene mediating the majority of trans genes (lower entropy) tend to yield more reproducible Rank-1 cis genes across bootstrapped resamples than SCNAs with multiple cis genes with similar trans mediation (higher entropy).

**Figure 4.**
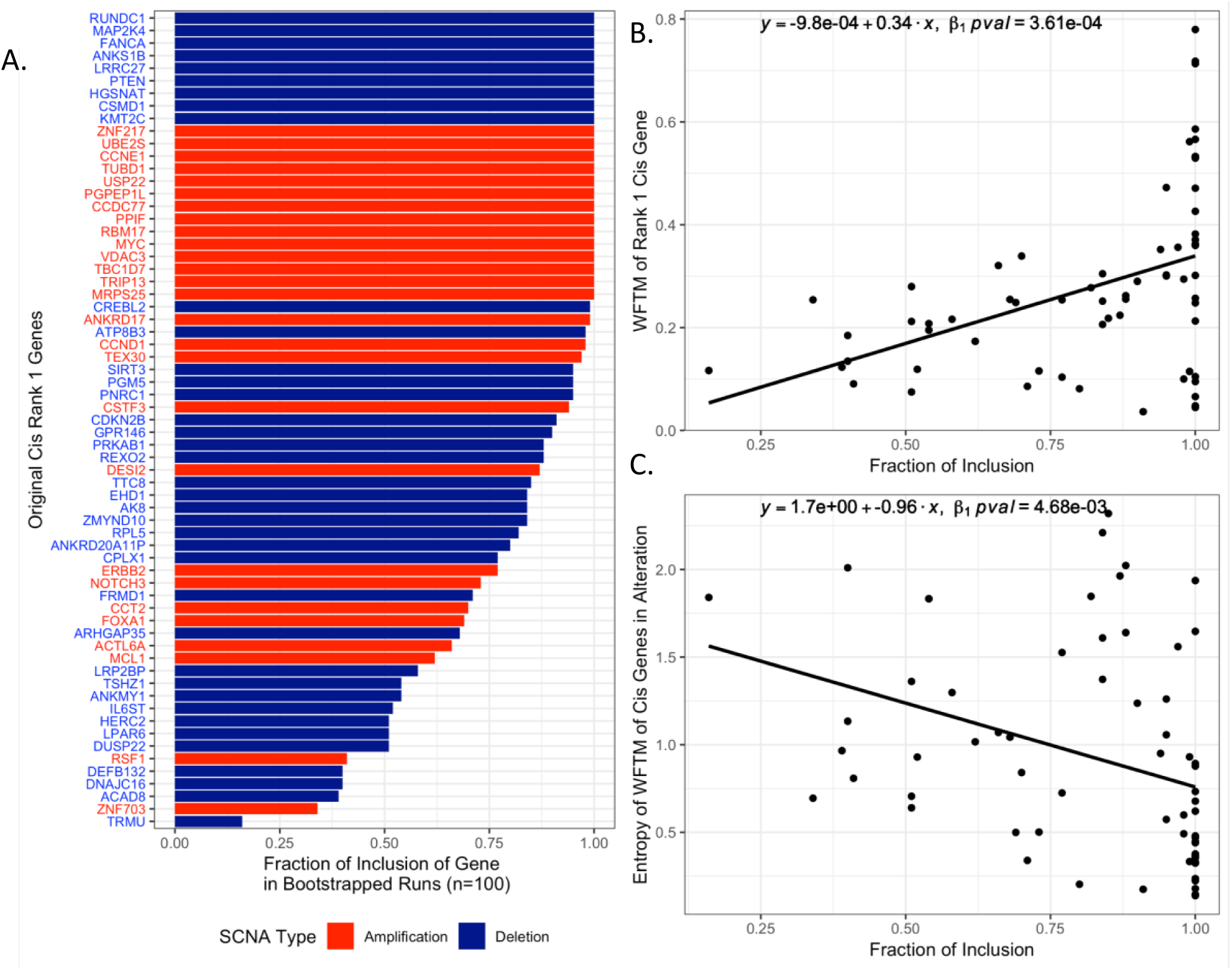
Reproducibility of Rank-1 cis genes across bootstrapped resamples. (A) Fraction (i.e., frequency) of inclusion of Rank-1 cis genes in bootstrapped resamples (B) Fraction of inclusion vs. Weighted Fraction of Trans Mediation (WFTM) (C) Fraction of inclusion vs. entropy of WFTM of cis genes in the SCNA

### Sensitivity and Specificity of mediation testing

Evaluation of the Sobel test of mediation was carried out using simulated data of true positives and true negative examples of mediation. Test performance, as measured using AUC, sensitivity and specificity, was recorded for varying combinations of correlation between the independent variable and mediator (“correlationXY”) and sample size (“N”) (Figure 5). High specificity (true negative rate) was consistently achieved for all input parameter ranges. Sensitivity (true positive rate) drops under conditions of low correlation and low sample size. Nevertheless, the conditions for lower sensitivity is not characteristic of real datasets that were used to test and validate iEDGE. Specifically, high correlation between SCNA and cis genes and between SCNA and trans genes is expected given that only cis and trans genes that were significantly expressed with respect to the SCNA were considered prior to mediation testing.

**Figure 5.**
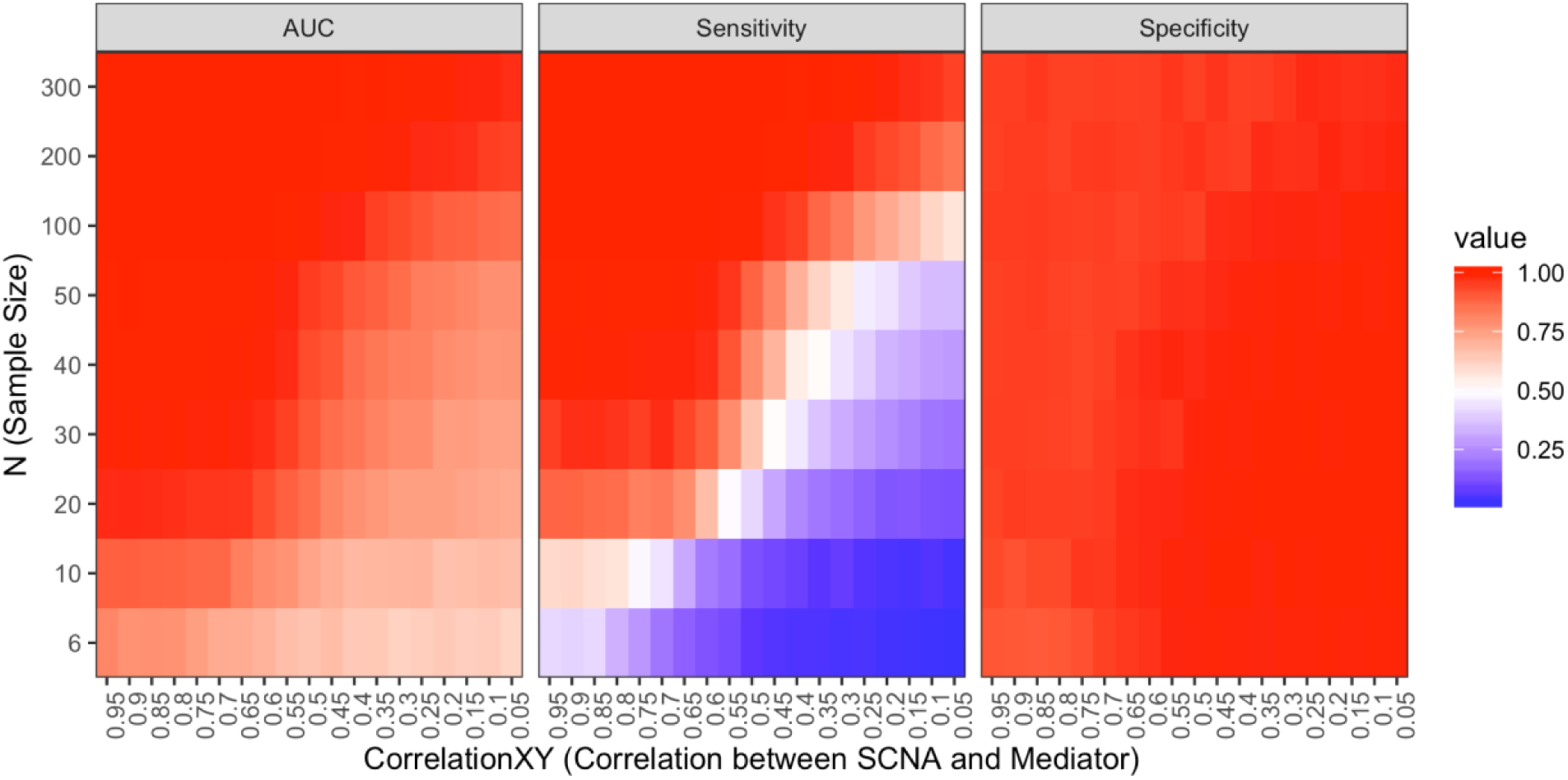
Mediation test performance on simulated data. AUC (Area Under the ROC Curve), sensitivity (true positive rate), specificity (true negative rate) for various values of correlationXY (correlation between SCNA and mediator, X-axis) and N (sample size, Y-axis).

Additionally, sufficient sample size is achieved in most of TCGA datasets tested (Supplemental Table S7), with the only exception being the TCGA Adrenocortical carcinoma dataset (n = 77) in which mediation results should be interpreted with caution.

### Graphical Portal of iEDGE Results Enable Targeted Queries

In order to enable fast interactive browsing of iEDGE precomputed runs on massive datasets such as the TCGA pan-cancer set, we developed a web portal (https://montilab.bu.edu/iEDGE/) to allow users to query iEDGE results selectively by cancer types, genes, and SCNAs. This portal displays graphical and tabular results for each step of the iEDGE pipeline, including differential expression of cis and trans genes, mediation analysis for driver prediction, and pathway enrichment analysis.

An overview of an example walkthrough of a targeted query is illustrated in Figure S2. Here, the TCGA breast cancer (BRCA) report is selected and the gene query is *ERBB2* (*HER2*). The table of differential expression results is available for the cytoband *17q12* in which *ERBB2* resides in. Additionally, results of the mediation testing and driver gene prediction is available in a bipartite graph format. In this case, the graph indicates that *ERBB2* is the top mediating cis gene and the predicted driver of the SCNA.

## Discussion

Methods developed for the integrative analysis of (epi-)DNA regulators and gene expression data often focus only on the genes harbored by the alteration regions, while not considering the downstream (trans-)effects, which may limit a method’s ability to detect cancer driver genes. We present a computational framework for the integrative analysis of (epi-)DNA and gene expression data for large-scale datasets, iEDGE, that is able to thoroughly catalogue the cis and trans effects of epi-(DNA) alterations, and to predict the most likely cis-driver genes based on the extent of their mediation of downstream trans-genes.

The first step of the iEDGE pipeline uses differential expression analysis to determine the cis and trans genes that are associated with the presence/absence of a particular epi-DNA alteration across samples. By measuring the alterations’ association with trans genes we capture meaningful biological mechanisms representing downstream effects that are generally missed by tools that only consider genes within the alterations (e.g., GISTIC2.0). Trans genes are of potential high relevance considering that (epi-)DNA regulators such as SCNAs harbor many upstream genes in signaling pathways, e.g., transcription factors, wherein the set of target genes effected can shed light on processes or pathways associated with disease progression.

The second step of the pipeline, the mediation analysis, ranks the set of cis genes by the extent of their mediation of trans genes. We showed that mediation analysis captures important cancer driver genes in our study of the TCGA breast cancer dataset. We then expanded these results by performing a pan-cancer analysis across 19 cancer types in the TCGA, further highlighting our tool’s ability to identify known, as well as potentially novel drivers.

We conducted extensive in silico validation of predicted cis driver genes, by first testing for their enrichment with known drivers from multiple cancer driver databases. We then characterized predicted drivers by testing for their “essentiality” against genetic screens and cellular model data included in the DepMap, to explore SCNA-associated gene dependencies. Both analyses showed that our list of predicted genes is significantly enriched for cancer genes of functional relevance, either as cancer drivers, potential cancer therapeutic targets, or markers of disease progression. Further experimental studies are needed to validate and characterize predicted drivers. Furthermore, simulations were performed to validate the performance of the Sobel test of mediation. Under assumptions of adequate sample size and strong correlation effects between SCNA and cis acting genes, conditions which were met in most of the TCGA datasets used in this study, the Sobel test of mediation achieved high AUC for uncovering true mediation effects.

Similar approaches have recognized the importance of integrating gene expression and the coordinated expression of affected gene modules for identifying drivers. One notable example is CONEXIC (Akavia et al., 2010), which identified functional gene modules for each candidate regulator in the form of copy number alteration (CNA). The conceptual steps of iEDGE are similar to the “Single Modulator Step” outlined in Akavia et al, in which, first, cis genes are defined in their “candidate driver gene” selection process, and trans genes are defined in their “target gene modules” selection step, and second, a scoring function is used to find the single candidate driver gene that best associates with the target gene expression module. The implementation details of the second step for the scoring of driver genes is different compared to iEDGE. iEDGE considers each 3-layer relationship between SCNA status, cis, and trans gene expression to calculate a mediation effect and to rank cis genes, whereas the “Single Modulator Step” of CONEXIC computes the best candidate driver using a Normal Gamma scoring function to measure each target gene’s fit with each candidate driver’s gene expression. Furthermore, while CONEXIC is not available for public use, iEDGE is available as an open-source R package (https://github.com/montilab/iEDGE) to enable analysis of custom datasets, and it supports the automatic generation of multi-level html reports in easily interpretable graphical formats as shown for each of the TCGA cancer types in the iEDGE web portal (montilab.bu.edu/iEDGE).

Other model-based methods incorporate models of causal relationships between multiple levels of genomic data through the scores of conditional dependencies, measured using partial correlation coefficients for normal continuous features (Amgalan and Lee 2015), or conditional mutual inclusive information for binary or mixed binary and continuous features (Kim et al. 2016; Zhang et al. 2014). These approaches are similar to the mediation testing step of iEDGE, but they are more conservative models that detect full mediation, e.g., significant hits are instances in which the conditional independence given the mediator is zero, a condition that is rarely satisfied by genomic data, whereas mediation tests are also able to capture partial mediation in which the association between the independent and response variable is significantly reduced in size when the mediator is introduced but may still be different from zero.

One assumption used in the iEDGE analysis presented in this study is that the number of trans genes that a cis gene mediates is a proxy measure of a gene’s importance (i.e., of its likelihood of being a cancer driver). This is a simple and intuitive heuristic to estimate the extent of transcriptional impact from each (epi-)DNA regulator, albeit it may not be an appropriate assumption for specific use cases. For more targeted analysis, one may be only interested in predicting the cis gene that mediates gene targets in a particular pathway. Our tool is customizable in that the user can specify the set of trans genes to consider for mediation based on their membership in a pathway of interest or an experimentally derived gene signature.

We use SCNA as an example of epi-DNA events to demonstrate a convenient use case for this tool as SCNAs can be used to easily define genomic boundaries for distinguishing cis and trans acting genes.

However, this tool can also be applicable to the integrative analysis of other genomic/epi-genomic data types such as DNA methylation, DNA mutations, and microRNA regulatory networks. Similarly, gene expression dataset can use a variety of quantitative gene-centric measures such as RNA-seq, microarray, or proteomic assays.

## Methods

### iEDGE overview

An overview of the iEDGE approach is summarized in Figure 1. Briefly, iEDGE integrates samples quantified from a gene expression profiling assay paired with one or more genomic or epi-genomic assays capturing information upstream of gene expression, such as SCNAs, DNA methylation, or microRNA expression. First, we identify the list of genes associated in “cis” with the upstream epi-DNA alteration. In the case of SCNAs, cis genes of each SCNA in this list are defined as genes within the focal peaks of the SCNA, and trans genes are defined as genes outside of the focal peaks. Then, iEDGE performs differential expression analysis to identify cis and trans genes significantly associated with each SCNA and, optionally, pathway enrichment analysis of each significant cis and trans gene sets (Figure 1A). Next, iEDGE predicts cis <u>driver</u> genes using mediation analysis, wherein each differentially expressed cis gene is ranked by the number of differentially expressed trans genes it mediates as determined using the Sobel test of mediation (Figure 1B).

### Copy number and gene expression pre-processing

We utilized the dataset of somatic copy number alterations and RNA-seq from the TCGA breast cancer cohort, which was preprocessed using Firehose v0.4.13 and downloaded from the Broad Institute TCGA GDAC repository (http://gdac.broadinstitute.org/runs/).

The SCNA dataset was preprocessed using GISTIC2.0 under the Firehose run release analyses 2015_08_21, which identified 29 significant focal amplifications and 40 significant focal deletions to be considered for integrative analysis (Broad Institute TCGA Genome Data Analysis Center 2015). SCNA status by sample was binarized using the amplitude threshold of 0.1, that is, SCNA status = 1: t < 0.1 or 0: t >= 0.1 for amplifications and 1: t< −0.1 or 0: t >= −0.1 for deletions. Cis genes were identified by GISTIC2.0 as genes in the wide peak of each significant SCNA with boundaries selected at the confidence level of 0.99. Trans genes were identified as genes outside the wide peak of each SCNA.

The gene expression data is a RSEM processed gene expression matrix (stddata 2015_06_01). Expression values were log_2_-transformed prior to integrative analysis. The samples were categorized into breast cancer subtypes using a combination of the pam50 classifier (Parker et al. 2009) and the *HER2* status. *HER2* status was determined using *HER2* receptor activity, labeled positive if tested positive by either FISH or IHC method. In *HER2* negative samples, the pam50 classification was used. Samples with gene expression-based membership in one of four major breast cancer subtypes (Luminal A, Luminal B, Her2, Basal) were retained for integrative analysis. Tumors classified as “Normal-like” by pam50 were removed from further analysis. A total of 1050 samples (primary solid tumors only) were found with paired gene expression and SCNA data by matching the sample barcode identifier (combination of patient id and sample type).

### Determining significantly expressed SCNA associated cis and trans genes

We performed differential expression of cis and trans genes with respect to each GISTIC2.0 defined significant focal SCNA peak. The significance of differential expression was estimated using limma (Ritchie et al. 2015) for each SCNA with samples split into two groups (amplified vs. normal for amplification peaks, and deleted vs. normal for deletion peaks) using FDR < 0.25 and fold change > 1.2 for cis genes, and FDR < 0.01 and fold change > 1.5 for trans genes. One-sided significant levels were reported for cis genes with the rationale that a focal amplification is commonly associated with an increase in gene expression and a deletion is associated with a decrease in gene expression. Two-sided significant levels were reported for trans genes, as indirect downstream effects can occur through either transcriptional repression or activation.

### Pathway enrichment analysis of significant cis and trans gene sets

Significantly differentially expressed cis and trans gene sets were tested for pathway enrichment using the MSigDB gene set compendia hallmark (hallmark gene sets), c2.cp (curated gene sets from online pathway databases), and c3 (motif gene sets), version 5.0 (Liberzon et al. 2011). The significance of pathway enrichment was determined using a hypergeometric distribution-based test and corrected for multiple hypothesis testing using the False Discovery Rate (FDR) method (Benjamini and Hochberg 1995).

For breast cancer-specific pathway enrichment results, pathways with significant enrichment (FDR < 0.25) in any SCNA were reported (Figure 2). In addition, each SCNA was labeled according to its over-representation in a particular breast cancer subtype using a one-sided Fisher exact test comparing counts of SCNA occurrence within vs. outside each breast cancer subtype (FDR < 0.05). Subtype-specific SCNAs were subsequently used in conjunction with pathway enrichment results to determine subtype-specific pathway enrichments using a one-sided Fisher exact test (FDR < 0.05).

### Mediation testing and prediction of cis drivers

To elucidate which cis genes are likely to mediate the association between copy number alteration and trans gene expression, we used the Sobel test to estimate the mediation effect of each cis gene and its significance (Sobel 1982). Briefly, we model the association for each triplet of SCNA, cis gene, and trans gene using the linear regression models specified in Figure 1B. The mediation effect of the cis gene: Δτ = τ - τ’, represents the change in the magnitude of the effect of the SCNA status on the trans gene expression after controlling for the cis gene expression. The significance of the mediation effect is calculated from the t statistic: t = Δτ /SE, where SE is the pooled standard error term, and is compared to the normal distribution to determine the p-value and FDR (Benjamini and Hochberg 1995).

An important simplifying model assumption is that the association between each SCNA and trans gene is mediated by at most one cis gene, therefore the cis gene mediator for each SCNA-trans gene pair is chosen based on the most significant mediation effect (ranked by the FDR values of the mediation test).

Once the cis genes mediator is determined for each unique trans gene, the mediation effect of cis gene *I* on trans gene *j* can be expressed as either binary (0 or 1), based on the significance of the mediation, or as the weight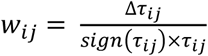, limited to the range of [0, 1]. Using the latter weighted approach, the total fractional weighted mediation effect of cis gene i across m significantly expressed trans genes, also referred to as the *Weighted Fraction of Trans Mediation* (WFTM), is expressed as 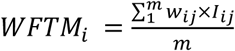 where I_ij_ denotes the indicator variable taking the value 1 if cis gene *i* is has the most significant mediation effect on trans gene *j* among all cis genes, 0 otherwise.

Next, for each SCNA, we rank each cis gene based on its total mediation effect M_i_. The cis gene with the highest value of M_i_ is denoted as the “Rank-1 cis gene” for the given SCNA, the candidate driver gene of the alteration.

### Assessing enrichment of predicted cis drivers in databases of known cancer drivers

To investigate the functional impact of putative drivers identified by iEDGE, we tested for the enrichment of iEDGE predicted driver genes in several cancer driver databases. Reference cancer driver genes, denoted as either “oncogenes” or “tumor suppressors” in the original sources, were compiled using data from Tuson Explorer (Davoli et al. 2013), Online Mendelian Inheritance in Man (OMIM) (Hamosh et al. 2005), Cancer Gene Census (CGC) from Catalogue of Somatic Mutations In Cancer (COSMIC) (“Cancer Gene Census” 2018; Forbes et al. 2017), and Uniprot (The UniProt Consortium, 2017). We tested for the overrepresentation among Rank-1 cis genes, compared to non-Rank-1 cis genes, of known drivers from the reference databases (Table S7). Enrichment tests were conducted separately for each reference database and driver type, i.e., oncogene (“_OG”), tumor suppressor (“_TN”), or both (“_COMBINED”), as well as using the union of the driver genes across databases (column “ANY” indicates union of drivers across knowledge bases). Enrichment significance was calculated using a one-sided Fisher exact test assessing the overrepresentation of Rank-1 vs. non-Rank-1 cis genes with respect to their membership in the reference driver list, conditional on the direction of change, e.g. amplified cis genes among oncogenes, deleted cis genes among tumor suppressors, or direction insensitive. P-values are adjusted with the FDR procedure to correct for multiple hypothesis testing across 19 tumor types.

### Copy number-associated gene dependencies

SCNAs often lead to overexpression of driver oncogenes and confer a tumor-promoting environment. In other words, driver oncogenes are more likely to act as essential genes (increased gene dependency) in an amplified state. To identity such genes, we looked for copy number associated gene dependencies using data available from DepMap, specifically, gene dependency data (McFarland et al. 2018) and cancer cell line genomics data from the Cancer Cell Line Encyclopedia (CCLE) (Barretina et al. 2012; Cancer Cell Line Encyclopedia Consortium 2015). In particular, we mined for genes with associations between gene dependency (Combined RNAi screens from Broad, Novartis, Marcotte) and somatic copy number status across cell lines (CCLE). To do this, we used a linear regression model Y = α X + β where Y is the Gene Dependency Score and X is the copy number level (log2 relative to ploidy) across cell lines. Gene dependency scores were calculated using DEMETER2 (McFarland et al. 2018). A negative gene dependency score corresponds to high gene dependency, e.g., an increased gene essentiality. In contrast, a high gene dependency score corresponds to non-essential genes. We looked for genes with significantly negative association between copy number and Gene Dependency Score, that is, genes in which higher copy number is associated with higher gene essentiality, using a one-sided t-test on the coefficient α (alternative hypothesis α < 0). Additionally, FDR correction was performed across p-values for all genes. Since gene dependency scores are calculated from only gene knockdowns, we were only able to test amplification-driven gene dependencies, whereas overexpression assays would be needed for detection of deletion-driven gene dependencies.

Finally, to determine if iEDGE was able to uncover an enrichment of amplification-driven gene dependencies, we tested for enrichment of genes with amplification-driven gene dependencies among iEDGE-predicted Rank-1 cis genes using a one-sided Fisher test on the contingency table of counts (rows: membership in Rank-1 vs. non-Rank-1, columns: presence vs. absence of amplification driven gene dependencies).

### Pan-Cancer Analysis

TCGA gene expression and copy number (GISTIC2.0) data were retrieved using Firehose v0.4.13 for 19 cancer types as summarized in Table S6. Gene expression data (RNASeq) correspond to the latest release at the time of retrieval (stddata 2015_02_04 for cancer types ACC, KIRP, THCA and stddata 2016_07_15 for all other cancer types). SCNA copy number data uses the GISTIC2.0 run corresponding to Firehose run release analyses 2016_01_28. Gene expression processing and copy number processing steps for the pan-cancer analysis are consistent with methods used for the BRCA-only analysis. Of note, the BRCA dataset in the pan-cancer analysis includes all TCGA BRCA samples with paired copy number and gene expression data to be consistent with processing of other TCGA cancer types, contrary to the removal of samples without an assigned molecular subtype in the BRCA-only analysis.

We tested for enrichment of known cancer driver genes among Rank-1 cis genes in each of the 19 TCGA cancer types using a one-sided Fisher test (Supplemental Table S8), consistent with the BRCA-only analysis. FDR correction was performed on the nominal p-values across all 19 cancer types for each test (unique combination of database origin and alteration direction, gain or loss). Enrichment tests are direction sensitive (“OG” tests for enrichment of oncogenes in Rank-1 cis genes in amplifications, “TN” tests for enrichment of tumor suppressors in Rank-1 cis genes in deletions, “COMBINED” tests for the union of the two sets). To determine the significance of the number of expected subtypes with increased sensitivity of driver predictions compared to random, we conducted a Kolmogorov-Smirnov test using the p-values of test of enrichments across subtypes against the uniform distribution in the range of [0,1].

We also tested for enrichment of amplification-driven gene dependencies in Rank-1 cis genes across the 19 cancer types (see Methods: Copy number-associated gene dependencies). Multiple hypothesis correction using the FDR procedure (Benjamini and Hochberg 1995) was used to adjust the significance values across multiple cancer types. Similarly, the significance of number of expected subtypes with increased sensitivity of amplification-driven gene dependencies is determined using a Kolmogorov-Smirnov test.

### Evaluation of the Reproducibility of Cis Driver Gene Predictions in BRCA

To evaluate the consistency of Rank-1 driver gene predictions, we generated 100 bootstrapped resamples of the original TCGA breast cancer dataset using sampling with replacement with the number of samples equal to the size of the original dataset, derived the predicted Rank-1 cis genes across the 100 bootstrapped datasets, and compared these predictions against the original list of predicted Rank-1 cis genes from the full dataset. A reproducibility score was calculated for each of the original predicted Rank-1 cis gene as the percent of inclusion of the particular gene as a Rank-1 cis gene among the bootstrapped results. To explain the variation on reproducibility scores across genes, we modeled these scores using linear regression models with the dependent variable being either the Weighted Fraction of Trans Mediated (WFTM) or the Entropy of WTFM for the alteration of interest, calculated as the Shannon Entropy of WTFM of all differentially expressed cis genes within the alteration harboring the Rank-1 cis gene of interest. A two-sided t-test on the slope, β_1_, of the linear regression, with H_a_: β_1_ ≠ 0, was used to estimate significance, defined as p-value < 0.05.

### Evaluation of Mediation Testing from Simulated Data

The Sobel test of mediation identifies cis genes that mediates SCNA and trans gene expression. To determine the conditions in which mediation is correctly identified, we used a forward simulation approach to generate labeled data of true positives and true negative, and then applied the mediation test to estimate its sensitivity and specificity. Of note, for simplicity, these simulated instances represent total, not partial, mediation.

True positive instances of mediation were generated using the following linear regression models:

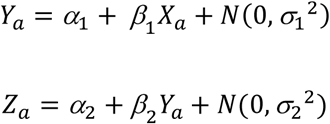

Here, X_a_ denotes the independent variable (SCNA status), Z_a_ is a dependent variable (trans gene expression) and Y_a_ is a true mediator of X_a_ and Z_a_ (cis gene expression).

True negative instances, representing the lack of a mediation effect, were generated using the following models:

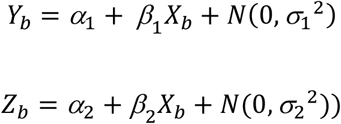

Here, X_b_ is the independent variable (SCNA status) and Y_b_ and Z_b_ are both dependent variables generated based on separate regression models from X_b_.

Variables X, Y, Z are vectors of length n, representing the sample size of the data. X is a binary vector (0s and 1s) corresponding to the binarized SCNA copy number status. σ is the standard deviation of the Gaussian noise term. We fixed β_1_ and β_2_ at 0.7 based on estimation from real data (TCGA breast cancer). The mediation test was performed on 1000 simulated true positives and 1000 simulated true negatives, and performance was measured in terms of AUC, sensitivity and specificity. For sensitivity and specificity, mediation calls were made based on the Sobel test p-value of 0.05. Test performance was recorded for simulated datasets based on a range of values of n (sample size) and σ (standard deviation of the Gaussian noise in the regression models). The standard deviation σ is a proxy for the correlation strength between dependent and independent variables, as higher noise corresponds to weaker correlation. For interpretability purposes, values for the parameter σ are converted to the corresponding Pearson correlation estimates using a Loess model (Local Regression).

### Software availability

iEDGE is available as an R package for download at https://github.com/montilab/iEDGE.

### Data Access

The datasets analyzed in this study are available from the Broad Institute TCGA GDAC (http://gdac.broadinstitute.org/runs/) as described in Methods. iEDGE reports on these datasets are available in an interactive web portal (https://montilab.bu.edu/iEDGE) to allow for exploration and mining of results in user-friendly tabular and graphical formats.

## Acknowledgements

We would like to thank TCGA and dbGap (phs000178.v9.p8) for granting access to RNA-seq and Copy Number Variation data. This work was funded in part by Find the Cause Breast Cancer Foundation (http://findthecausebcf.org/), Boston University Superfund Research Program (P42 ES007381-21), and the National Institute on Aging (NIA cooperative agreements U19-AG023122, and R01AG061844).

