## supplemental materials for "iEDGE: integration of Epi-DNA and Gene Expression and applications to the discovery of somatic copy number-associated drivers in cancer"

**Table of Contents**

**1. Supplemental Tables (see supplementaltables.xlsx)**

***Table S1: Differential Expression of Cis Genes in Somatic Copy Number Alterations (SCNA) in TCGA Breast Cancer***

***Table S2: Differential Expression of Trans Genes in Somatic Copy Number Alterations (SCNA) in TCGA Breast Cancer***

***Table S3: Cis Genes Ranking Using Mediation Analysis for TCGA Breast Cancer***

***Table S4: Rank 1 Cis Genes for TCGA Breast Cancer***

***Table S5: Enrichment of Rank 1 Cis Genes in Cancer Driver Databases in TCGA breast cancer***

***Table S6: Amplification-driven gene dependencies among cis genes in amplifications in TCGA Breast Cancer***

***Table S7: Summary of TCGA datasets used in iEDGE pancancer analysis***

***Table S8: Enrichment of Rank 1 Cis Genes in Cancer Driver Databases in TCGA pancancer analysis (19 cancer types)***

***Table S9: Enrichment of Amplification-driven gene dependencies among rank 1 cis genes in amplifications in TCGA pancancer analysis (19 cancer types)***

***Table S10: Rank 1 cis genes ordered by number of occurrence across cancer types***

***Table S11: Top differentially expressed (D.E.) cis gene in each SCNA ordered by number of occurrence across cancer types***

***Table S12: All differentially expressed cis genes in SCNAs ordered by number of occurrence across cancer types***

**2. Supplemental Figures**

***Figure S1.*** Hierarchical clustering of somatic copy number alteration status (SCNA) across subtyped TCGA breast cancer samples

SCNA_Status legend key: -2 (deep loss, possibly homozygous deletion): t > -0.9; -1 (shallow loss, possibly heterozygous deletion): -0.9 ≤ t < -0.1; 0 (normal, diploid): -0.1 ≤ t < 0.1; 1 (low-level amplification): 0.1 ≥ t > 0.9; 2 (high-level amplification): t > 0.9

***Figure S2.*** iEDGE web portal overview

(A) Selection of iEDGE report by cancer type

(B) Selection of query gene

(C) Differential expression table report for cis genes

(D) Graphical report of mediation testing and driver prediction
